# Determination of Copy Number Variations and Affected Gene Networks in Breast Cancer from Mexican Patients

**DOI:** 10.1101/2025.08.28.672450

**Authors:** Violeta Larios-Serrato, Hilda-Alicia Valdez-Salazar, Javier Torres, Margarita Camorlinga-Ponce, Patricia Piña-Sánchez, Héctor Mayani, Martha-Eugenia Ruiz-Tachiquín

**Affiliations:** Laboratorio de Biotecnología y Bioinformática Genómica, Escuela Nacional de Ciencias Biológicas (ENCB), Instituto Politécnico Nacional (IPN), Unidad Profesional Lázaro Cárdenas, Prolongación de Carpio y Plan de Ayala S/N. Col Santo Tomás, Alcaldía Miguel Hidalgo, CP 11340, Ciudad de México, México; Unidad de Investigación Médica en Enfermedades Infecciosas y Parasitarias (UIMEIP), Unidad Médica de Alta Especialidad (UMAE)-Hospital de Pediatría “Dr. Silvestre Frenk Freund”, Centro Médico Nacional Siglo XXI, Instituto Mexicano del Seguro Social (IMSS). Av. Cuauhtémoc 330, Colonia Doctores, Alcaldía Cuauhtémoc, CP 06720, Ciudad de México, México; Unidad de Investigación Médica en Enfermedades Oncológicas (UIMEO), Unidad Médica de Alta Especialidad (UMAE)-Hospital de Oncología, Centro Médico Nacional Siglo XXI, Instituto Mexicano del Seguro Social (IMSS). Av. Cuauhtémoc 330, Colonia Doctores, Alcaldía Cuauhtémoc, CP 06720, Ciudad de México, México

**Keywords:** CNV, genes, interaction networks, breast cancer, high-density arrays

## Abstract

Triple-negative breast cancer (TNBC) is an aggressive subtype with limited treatment options and high molecular heterogeneity. In this study, we performed a genome-wide analysis of copy number variations (CNVs) using high-density microarrays in tumor tissue (TUM), tumor adjacent tissue (ADJ), and leukocytes (LEU) from five Mexican TNBC patients. We identified both unique and shared CNVs across tissues, including alterations in key chromosomal regions such as 1q23.3, 1q32.1, and 8q24.3, which harbor oncogenes like MYC, MCL1, and BCL9. Losses in 6q25.2 affecting ESR1 were also detected. CNVs were enriched in genes related to the Hallmarks of Cancer, with TUM samples showing profiles associated with proliferation, metastasis, and immune evasion; ADJ samples with growth suppression; and LEU samples with genomic instability. Pathway enrichment analyses revealed disrupted functions in DNA repair, extracellular matrix organization, and TP53 signaling in TUM. Notably, EGFR, ERCC4, and HSP90AB1 emerged as central nodes in interaction networks and may serve as markers or therapeutic targets. This is the first CNV profiling study of its kind in TNBC from Mexican patients, highlighting the importance of including underrepresented populations in genomic research to uncover distinct molecular signatures and potential diagnostic or therapeutic avenues.

**Implications:** Molecular signatures of breast cancer (BC), predicted bioinformatically, involve common and distinct CNV-Hallmarks of Cancer genes, which are suitable candidates for screening as potential BC markers.

## Introduction

Despite advances in our understanding of cancer biology, breast cancer has the highest incidence (9,664,889 new cases) and mortality (4,313,548 deaths) in women Worldwide (Globocan, 2022, accessed March 21, 2025).

Cancer is a group of genetic diseases, thus creating the need to continue generating knowledge that helps us determine markers for this neoplasia to prevent, diagnose, predict response to treatments, and estimate survival. Based on the above, our interest focused on the analysis of copy number variations (CNVs), specifically the determination of changes in the number of copies of a particular DNA sequence in an individual’s genome. CNVs include insertions, deletions, and duplications of DNA segments, and explain a significant proportion of the genetic variability between individuals and cancer patients. Analyzing these alterations in breast cancer (BC) patients globally to generate sets of altered genes (losses or gains), they are grouped into signatures or profiles, and these into interaction networks that land on the Hallmarks of Cancer, will help find new marker molecules for neoplasia in the different molecular subtypes. We focused our interest on samples from patients with negative hormonal markers and HER2, that is, in the so-called triple-negative breast cancers (TNBC), where treatment options are limited, and tumor aggressiveness is higher. Seeking to provide a personalized and/or precision medicine approach, we were interested in analyzing the tumors, the adjacent tissue of the tumors, and the peripheral blood of each patient [estrogen and progesterone negative receptors (ER- and PR-), and human epidermal growth factor receptor 2 negative (HER2-)].

BC is the most common cancer among women in developed countries, with a significant contribution from genetic susceptibility factors. Despite extensive research, a substantial proportion of hereditary-BC predisposition remains unexplained, indicating the need for further exploration of genetic variations.

Recent studies have identified CNVs as prevalent structural variations in the Human genome, contributing to the risk of developing BC (Krepischi et al., 2012). CNVs involve alterations in the number of copies of specific DNA segments and can range from small alterations to significant chromosomal changes. While small CNVs are typically benign, larger ones can be linked to severe consequences such as developmental disorders and cancers (Hernández-Gómez et al., 2022). High-resolution genome-wide scans have demonstrated that BC patients show a higher frequency of rare CNVs compared to healthy controls, suggesting that these variations play a critical role in hereditary-BC risk (Pylkäs et al., 2012). The complexity of BC is further highlighted by its molecular heterogeneity, which is categorized into several subtypes based on specific genetic and phenotypic features (Krepischi et al., 2012). These subtypes include Luminal A, Luminal B, HER2+, and Basal types, each characterized by distinct receptor status and responses to treatment (Krepischi et al., 2012). Notably, CNVs associated with these subtypes have been shown to influence gene expression, molecular subtyping, and personalized treatment strategies, emphasizing the importance of understanding CNVs in the context of BC (Hernández-Gómez et al., 2022). As research advances, the potential for CNV profiles to serve as effective biomarkers for BC diagnosis and prognosis continues to grow (Hernández-Gómez et al., 2022). In the context of Latin America, founder mutations have been identified, highlighting the ethnic diversity in genetic variations affecting BC risk. Understanding these variations and their associated gene networks in Mexican patients is essential for developing targeted therapies and improving clinical outcomes.

## Materials and Methods

### Samples

The present study included five adult females (CM09, CM10, CM27, CM30, and CM64), see Table 1. The pathological characteristics of the samples were determined by two independent pathologists.

**Table 1.** Patient and sample characteristics.

| ID | Sample type | Cancer type | Gender | Age (years old) | TNM | Clinical stage | Tx | % of cancer cell |
| --- | --- | --- | --- | --- | --- | --- | --- | --- |
| CM09 | LEU<br>ADJ<br>TUM | IDC | F | 80 | T2N0M0 | IIA | Without | 95 |
| CM10 | LEU<br>ADJ<br>TUM | IDC | F | 35 | T2N2M0 | IIIA | Without | 100 |
| CM27 | LEU<br>ADJ<br>TUM | LC | F | 88 | T2N1M0 | IIB | Without | 80 |
| CM30 | LEU<br>ADJ<br>TUM | IDC | F | 64 | NA | NA | Without | 70 |
| CM64 | ADJ<br>TUM | IDC | F | 61 | T2N0M0 | IIA | Without | 70 |

ID: Identification code; LEU: leukocytes; ADJ: tumor adjacent tissue, and TUM: tumor tissue; infiltrating ductal carcinoma: IDC; lobular carcinoma: LC; F: female; TNM: tumor size, ganglion numbers, metastasis classification; Tx: treatment; NA: not available.

The study inclusion criterion was individuals who attended the Oncology Hospital of the XXI Century National Medical Center-Mexican Institute of Social Security (IMSS), Mexico City, Mexico. The study exclusion criteria were individuals and samples without a TNBC diagnosis, patients with other cancer types, and patients with recurrent cancer. Samples were excluded from the study if their DNA quantity and/or quality were insufficient for analysis.

The present study was approved (R-2017-3603-65) by the Scientific and Ethics Committees of the Mexican Institute of Social Security (IMSS). The study participants were informed of its nature and asked to sign a consent form. The study was conducted according to the best clinical practices of our institution. The identities of all study participants were anonymized.

### Immunohistochemistry

The data were obtained from the pathology and clinical records of the patients (Table 2). All samples were triple-negative breast cancer (TNBC).

**Table 2.** Immunohistochemistry data samples.

| ID | ER | PR | HER2 | Ki67 | CK5 |
| --- | --- | --- | --- | --- | --- |
| CM09 | Negative | Negative | Negative | Negative | Positive |
| CM10 | Negative | Negative | Negative | Negative | Negative |
| CM27 | Negative | Negative | Negative | Positive | Negative |
| CM30 | Negative | Negative | Negative | Negative | Negative |
| CM64 | Negative | Negative | Negative | Positive | Positive |

ID: Identification code; ER: Estrogen receptor; PR: Progesterone receptor; HER2: Human epidermal growth factor receptor 2; Ki67: Marker of proliferation Ki-67; CK5: Cytokeratin 5.

### DNA extraction

DNA extraction was done with a commercial kit (QIAamp® micro-Kit, QIAGEN®) according to the manufacturer’s instructions. The extraction was modified to include an initial incubation at 95°C for 15 min, followed by 5 min at room temperature as described previously, before being digested with proteinase K for three days at 56°C in a water bath, adding fresh enzyme at 24 h intervals.

### DNA quality assessment and preparation

The extracted DNA was quantified by spectrophotometry (Nanodrop 2000, Thermo Scientific). Multiplex PCR was done to assess the quality of DNA (Multiplex PCR kit, QIAGEN®) with a set of primers to amplify various regions of the GAPDH gene (Utrera-Barillas et al., 2013). Products were visualized by electrophoresis (RedGel® Nucleic acid gel stain, Biotium) on a 1% agarose gel and documented under an ultraviolet light transilluminator system (Syngene, Frederick, MD, USA).

### High-density whole-genome microarray analysis

Samples were analyzed by Affymetrix® CytoScan™ microarrays according to the manufacturer’s protocol, beginning with 250 ng DNA, except for the addition of five PCR cycles to increase the DNA sample. PCR products (90 micrograms) were fragmented and labeled using an additional PCR.

### Copy number processing

Raw intensity files (.CEL) retrieved from the commercial platform were analyzed using their proprietary software Chromosome Analysis Suite v4.3.0.71, using the CytoScanHD_Array.na36.annot.db file for annotation, and the GRCh38 genome (February 2013) as a reference model.

Data processing was based on the segmentation algorithm, where the Log2 ratio for each marker was calculated relative to the reference signal profile. To calculate CNVs, the data were normalized to baseline reference intensities using the reference model (provided by ChAS), including 284 HapMap samples, as well as 96 healthy normal individuals. The Hidden Markov Model (HMM) available in ChAS was used to determine the copy number state (CN-state) and their breakpoints. The customized high-resolution condition was used as a filter for the determination of CNVs: CN-gains with 50 marker count and 400 Kbp, and CN-losses with 50 marker count and 100 Kbp. We used the median absolute pairwise difference (MAPD) and the single-nucleotide polymorphism quality control (SNP-QC) score as the quality control parameters. Only samples with values of MAPD < 0.25 and SNP-QC > 15 were included in further analysis.

### Bioinformatic analysis

We developed a custom Perl script to process the CNV segment data files generated by ChAS for each sample, compare the files to build a table of genes that contains frequencies in patients, events types (gains or losses), chromosomes and cytogenetic bands, and Online Mendelian Inheritance in Man (OMIM) information, and incorporate additional information from different databases: haploinsufficiency information from DECIPHER (Firth et al., 2009) database of genomic variation, genes reported at dbEMT 2.0v (Zhao et al., 2019) (Table Supplementary 1).

The genes altered in at least three patients (cut-off ≥ 3) with BC were included for analysis and visualizations with the R language v4.4.0 and RStudio 2025.05.0+496 environment for statistical computing and graphics, with packages from Bioconductor v3.21.

The genes altered in at least three patients (cut-off ≥ 3) in TUM, ADJ, and LEU were included for analysis and visualization. A karyotype was built with KaryoploteR 1.30.0 and biomaRt 2.60.1. The genes were compared by generating Venn diagrams with the Jvenn server (Bardou et al. 2014). Cancer Hallmarks enrichment analysis (*p*.adjust < 0.05) was performed with a collection of 6,763 genes (Menyhart et al., 2024). Reactome v88 performed a metabolic pathway enrichment analysis (Fabregat et al., 2018), considering those results significant with values less than 0.05 in the false discovery rate (FDR). Finally, to establish the profile-associated Hallmarks of Cancer involving TUM, ADJ, and LEU, we generated an interaction network by CNV-type (gains and losses) based on genetic and physical interactions, biological pathways, and predicted relationships using the String, GeneMANIA (Warde-Farley et al., 2010) prediction server and Cytoscape v.3.10.3 (Shannon et al., 2003), including the manual annotation of their corresponding Cancer Hallmarks: adhesion, angiogenesis, inflammation, migration, metastasis, morphogenesis, proliferation, and survival (Hanahan and Weinberg 2011) with punctual scrutiny and help from databases such as The Human Protein Atlas (Uhlen et al., 2017).

## Results

### Sample characteristics

Samples from five patients with BC from Mexico (third-generation Mexicans) between 80 and 35 years of age (mean ± SD, 65.6 ± 20.42 years), four without any previous cancer (naïve), and one with treatment were included in the present study.

Each sample has TUM, ADJ, and LEU, except CM64 (a total of 14 samples were analyzed). The raw data were deposited in the NCBI Gene Expression Omnibus database (ID GSE290707).

Table 1 presents the ID, cancer type, four samples with infiltrating ductal carcinoma (IDC), and one with lobular carcinoma (LC); three samples with clinical stage IIA, one with IIIA, and one without data on the medical record, and the percentage of neoplastic cells for tumor tissues ranging between 70 and 100%. All these are issues about the complexity of genetic analysis. In this study, we analyzed samples from five female patients (CM09, CM10, CM27, CM30, and CM64) diagnosed with BC, including both IDC and LC. The samples consisted of TUM, ADJ, and LEU. The patients ranged in age from 35 to 88 years old. Clinical staging included T2N0M0 (stage IIA) for CM09 and CM64, T2N2M0 (stage IIIA) for CM10, T2N1M0 (stage IIB) for CM27, and no available data for CM30. All patients except CM64 were not receiving treatment at the time of sample collection. Immune cell infiltration levels, as measured, ranged from 70 to 100 across the cases.

The immunohistochemical analysis of the samples revealed a heterogeneous profile across the panel of biomarkers, as seen in Table 2. All five cases (CM09, CM10, CM27, CM30, CM64) were negative for HER2 expression. Estrogen receptor (ER) expression was consistently negative. Progesterone receptor (PR) was negative in all samples. Ki67 expression was positive in CM27 and CM64, suggesting a higher proliferative index in these tumors. Interestingly, CK5 was positive in two cases (CM09 and CM64), indicating a possible basal phenotype. Overall, the cohort displayed a predominance of TNBC features, with variability in proliferation and basal markers.

### Genomic detection of CNVs

The CNVs of the patients were estimated using the analysis described before, a meticulous process based on regions where the preponderance of SNPs does not display heterozygosity. Table 3 shows a summary of the chromosomes with the highest involvement frequency concerning the number of events (coincidences) at the CNVs, but not strictly perfect in the chromosomal coordinates.

**Table 3.** Principal affected chromosomes by CNVs-cumulative length in breast cancer tumor tissues, adjacent tissues, and leukocytes.

| Sample type | Chromosome | Number of events-GAINS | Number of events-LOSSES | Cumulative length-GAINS, Mbp-cl | Cumulative length-LOSSES, Mbp-cl |
| --- | --- | --- | --- | --- | --- |
| TUM | 1 | 77 | 3 | 64.82 | 2.38 |
|  | 3 | 129 | 8 | 71.16 | 3.41 |
|  | 5 | 16 | 32 | 21.52 | 97.18 |
|  | 7 | 35 | 23 | 56.93 | 27.07 |
|  | 13 | 1 | 18 | 12.85 | 64.94 |
| ADJ | 1 | 10 | 0 | 3.87 | 0.00 |
|  | 2 | 2 | 0 | 4.75 | 0.00 |
|  | 4 | 4 | 1 | 1.19 | 0.42 |
|  | 8 | 19 | 0 | 9.04 | 0.00 |
|  | 20 | 18 | 0 | 5.30 | 0.00 |
| LEU | 14 | 4 | 4 | 1.77 | 2.44 |

TUM: tumor tissues; ADJ: tumor adjacent tissues; LEU: leukocytes; Mbp-cl: megabase pairs-cumulative length.

Our data, which includes the megabase pairs cumulative length (Mbp-cl) of our tissue samples, were also reviewed (Table Supplementary 1). The CNV-gene frequency data, chromosomes, and cytobands are presented. Table Supplementary 2 displays the accumulated CNV-length (Mbp) values per chromosome to determine if more extended losses indicate more damage. To identify the most relevant CNVs in BC, we analyzed alterations occurring in at least three patients (cut-off ≥ 3). We found a similar pattern for total CNVs, with more events in the TUM (1717), ADJ (132), and LEU (13) (Table Supplementary 1).

In TUM, the affected chromosomes with Mbp-cl and the specified number of CNV-events were 3 (gains) and 5 (losses); at ADJ, they were chromosomes 8 (gains) and 4 (losses); LEU, chromosome 4 (gains and losses), see Table Supplementary 2. Interestingly, in TUM and ADJ, the most frequent CNV lengths were 200-500 Kbp (gains) and 100-200 Kbp (losses), see Table Supplementary 3.

Table 4 summarizes CNVs detected in Mexican BC patients across three tissue types: TUM, ADJ, and LEU (full information in Table Supplementary 4). In TUM, recurrent gains were observed in 1q23.3 (12.77 Mbp-cl) and 1q32.1 (7.03 Mbp-cl), as well as in the 8q24.22–8q24.3 region (20.32 and 16.25 Mbp-cl, respectively), while a loss was noted in 6q25.2 (3.18 Mbp-cl). ADJ displayed gains in cytobands 1p36.32, 4p16.1, 5p15.33, 8q24.3, and 20q13.33. LEU presented both gains and losses, including a loss at 8p11.22 and 14q11.2, and gains at 14q32.33 and 22q11.22.

**Table 4.** Cytobands and chromosomes affected by CNVs-cumulative length in tumor tissues, tumor adjacent tissues, and leukocytes.

| Type | Chromosome | Cytoband | Length<br>(Mbp-cl) | Type | Number of<br>patients |
| --- | --- | --- | --- | --- | --- |
| TUM | 1 | q23.3 | 12.77 | Gain | 5 |
|  | 1 | q32.1 | 7.03 | Gain | 5 |
|  | 6 | q25.2 | 3.18 | Loss | 5 |
|  | 8 | q24.22 | 20.32 | Gain | 5 |
|  | 8 | q24.3 | 16.25 | Gain | 5 |
| ADJ | 1 | p36.32 | 3.61 | Gain | 5 |
|  | 4 | p16.1 | 3.84 | Gain | 5 |
|  | 5 | p15.33 | 5.15 | Gain | 4 |
|  | 8 | q24.3 | 9.03 | Gain | 5 |
|  | 20 | q13.33 | 4.98 | Gain | 5 |
| LEU | 8 | p11.22 | 0.47 | Loss | 3 |
|  | 14 | q11.2 | 1.77 | Loss | 4 |
|  | 14 | q32.33 | 2.44 | Gain | 4 |
|  | 22 | q11.22 | 0.77 | Gain | 3 |

TUM: tumor tissues; ADJ: tumor adjacent tissues; LEU: leukocytes; Mbp-cl: megabase pairs-cumulative length.

Several of these cytobands have been frequently reported in BC. Gains in 8q24.3 and 20q13.33 are associated with amplification of oncogenes such as MYC and ZNF217, respectively, which promote tumor proliferation and metastasis (Curtis et al., 2012; TCGA Network, 2012). Likewise, gains on chromosome 1q—observed here at 1q23.3 and 1q32.1—are well-known markers of poor prognosis in BC and have been linked to the overexpression of genes such as MCL1 and BCL9 (Chial, 2008). The loss at 6q25.2 is particularly relevant, as it encompasses the ESR1 gene, which encodes the estrogen receptor frequently disrupted in hormone-sensitive BC (Curtis et al., 2012).

These findings suggest that CNVs are not limited to TUM but also occur in ADJ and LEU, which may reflect early or systemic genomic instability. Such alterations could have potential applications as markers or offer insights into the interaction between the tumor and the microenvironment.

### Breast cancer genes are associated with CNVs

A Venn diagram was constructed to examine the CNV-BC-relevant genes of at least three patients (cut-off ≥ 3) of the TUM, ADJ, and LEU (Figure 2A). We determined 1615 TUM, 25 ADJ, and one LEU CNV-uniques-genes (Figure 2 and Table Supplementary 5). From each subset (Figure 2D), those genes with matches according to the Cancer Hallmarks Genes database, a comprehensive resource that includes 133 genes, are shown, non CNV-genes were found in LEU; ADJ had 25 affected unique genes, and 100 in TUM. The six common genes TUM, and ADJ present a greater number of Hallmarks than LEU, which showed fewer. Figure 2. B-D represents the enrichment of the Hallmarks of Cancer.

**Figure 1.**
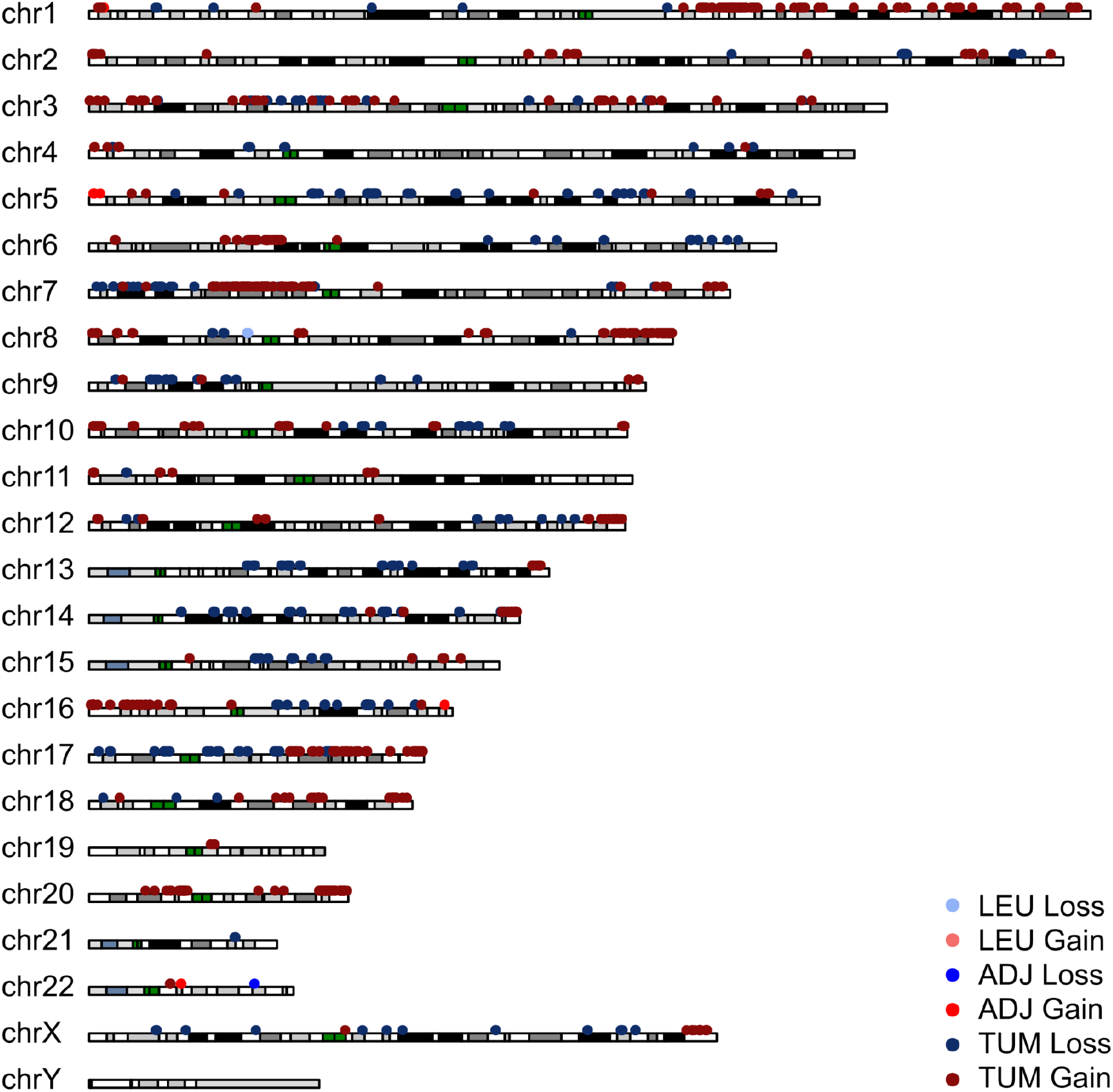
Karyogram with CNVs distribution in breast cancer samples. CNV events (gains or losses) were present from chromosomes 1 to 22, X, and Y. Gains (reddish colors) and losses (blue colors) are plotted for ≥ 3 tumor tissue (TUM), tumor adjacent tissue (ADJ), and leukocyte (LEU) samples. Cytobands (gray, black, or white bars) and centromeres (green bars) are shown.

**Figure 2.**
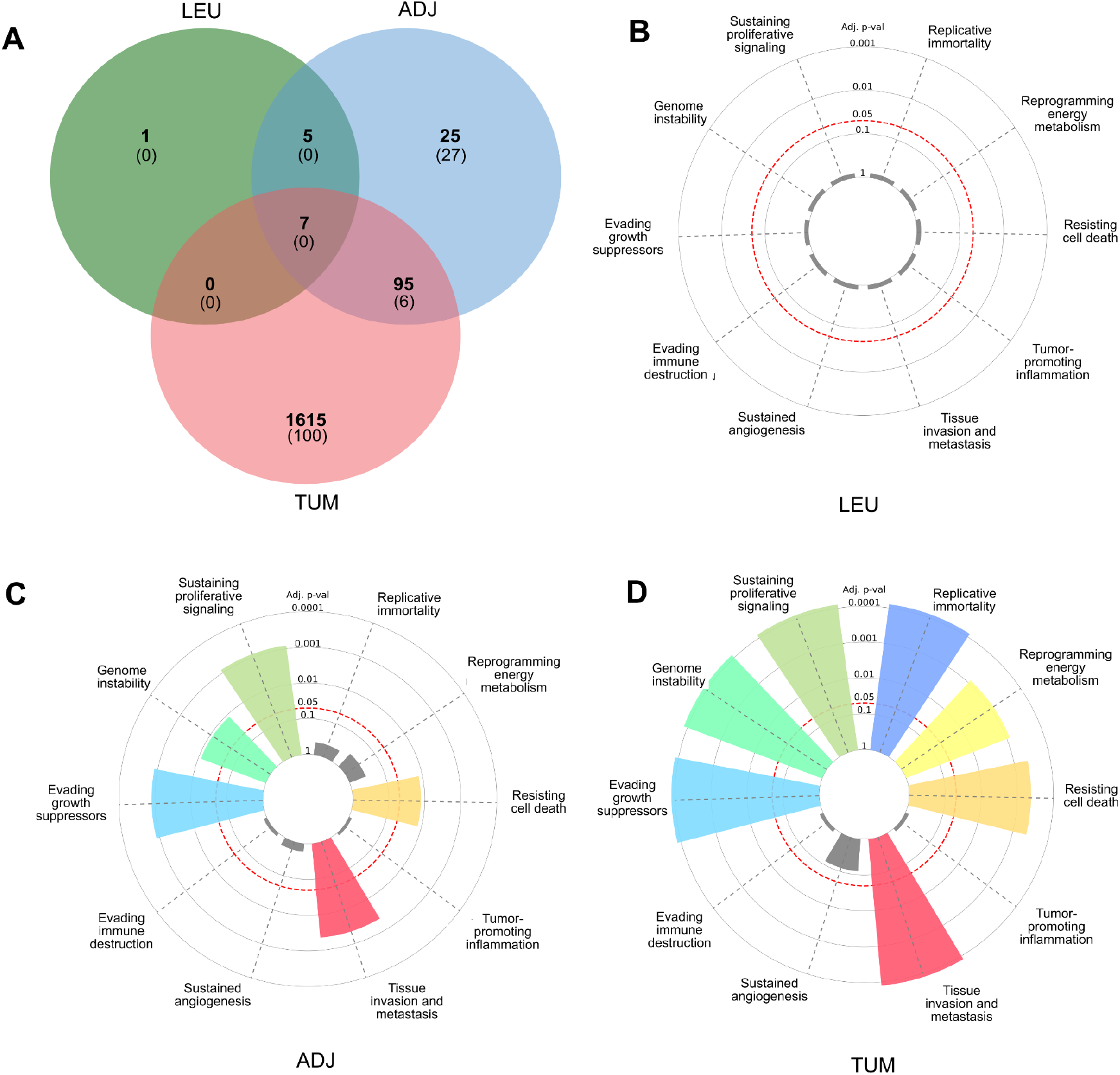
The Hallmark of Cancer enrichment plot. Profile of CNV-genes in breast cancer from ≥ 3 patients. The red dotted line represents the significance cut-off value (*Adj. p*.*val* < 0.05). Figure 2-A, the Venn diagram presents frequencies of specific and shared genes in tumor tissue (TUM), tumor adjacent tissue (ADJ), or leukocytes (LEU) in CNVs. Each subset of the Venn diagram in parentheses indicates the Cancer Hallmark genes. Figures B-D. In the radial graphs, each bar represents a Hallmark, and the height of the *p*.adjust value of the category enrichment; the grays have no significance.

The panels present Hallmark of Cancer enrichment plots, which visualize the statistical significance processes. Each radial plot depicts categories (e.g., genome instability, resisting cell death, tissue invasion, and metastasis), their bar lengths corresponding to adjusted *p-values*. Red dashed lines indicate common significance thresholds (0.1, 0.05, 0.01, and 0.001). Notably, the enrichment profiles vary across conditions, suggesting differential activation of cancer-related pathways. For instance, the top right plot shows strong enrichment in tissue invasion and metastasis. In contrast, the bottom right panel indicates minimal enrichment, suggesting a lack of Hallmark activation in that dataset. These comparative insights provide a functional context for CNV-genes and support mechanistic interpretations related to tumor progression and heterogeneity.

The genes altered in TUM are involved in sustaining proliferative signaling, tumor-promoting inflammation, tissue invasion and metastasis, evading immune destruction, resisting cell death, evading growth suppressors, reprogramming energy suppressors; ADJ, evading growth suppressor, and leukocytes, genome instability, and others, but no significance.

### Functional pathway analysis

Using the CNV-genes-Hallmarks of Cancer identified in each subset, metabolic pathways were predicted in Reactome using *Homo sapiens* as a model organism; in Table 5, those significant metabolic pathways (*p*-value < 0.05) and a brief general description are reported.

**Table 5.** Metabolic pathways and genes of the Hallmarks of Cancer.

|  | General description | Pathway | *p-value* | Genes |
| --- | --- | --- | --- | --- |
| TUM | DNA repair | Dual incision in TC-NER and GG-NER, TP53 regulates transcription | 9.5E-03 | CDK7, ERCC4, UVSSA, POLE2, POLD2, GTF2H2, ERCC6, CDK12 |
|  | Extracellular matrix/Collagen | Laminin interactions, assembly of collagen fibrils, collagen degradation | 1.6E-04 | COL4A1, COL4A2, ITGA3, LAMB3, PRSS2 |
|  | Cell signaling | Activation of TRKA receptors, signaling to RAS, | 7.7E-03 | ALK, EGFR, NTRK1, PIK3CB, RALA, |
|  |  | ERKs, PI3K |  | ADCYAP1R1, NRG2 |
|  | Cell cycle and checkpoints | G2/M checkpoints, DNA replication checkpoint, BRCA2, ATR | 2.1E-02 | CCNB1, CCNB2, HUS1, RAD17 |
|  | Regulation by TP53 | Formation of the early elongation complex, transcription of death receptors and ligands | 3.6E-02 | CDK12, CDK7, IGFBP3, PMS2 |
|  | Development and diseases | Chondrocyte maturation, defective LFNG causes SCDO3, disorders of developmental biology | 2.9E-02 | GLI2, HDAC4, NCOR2, NOTCH1, TBL1XR1 |
|  | Cellular stress response | HSF1-dependent transactivation, attenuation phase | 1.0E-03 | HSF1, CAMK2B, HSP90AB1, HSPA6 |
| ADJ | Signal transduction | Signaling NOTCH2, RAC3 GTPase cycle | 1.8E-03 | EP300, HES5, ARHGAP39, BAIAP2 |
|  | Neuronal System | Sodium permeable postsynaptic acetylcholine nicotinic receptors | 2.0E-02 | CHRNA4 |
|  | Metabolism | Alanine metabolism | 2.0E-02 | GPT |
|  | Developmental Biology | MITF-M-dependent genes involved in DNA replication, damage repair and senescence, transcriptional factor binding | 2.3E-02 | TERT, SOX8, EP300 BAIAP2, TP73, PRDM16 |
|  | Immune system | Co-inhibition by BTLA | 3.3E-02 | TNFRSF14 |
| LEU | Metabolism and antigen binding | Biological oxidations | 1.2E-01 | IGLL5 |
|  |  | B cell activation | 3.6E-02 |  |

TUM: Tumor tissues; ADJ: tumor adjacent tissues, and LEU: leukocytes.

### Correlation gene networks

Cancer CNV-genes-Hallmarks of Cancer associated with metabolic pathways were used to construct interaction networks by String (Figure 3). The connecting lines indicate associations by metabolic pathways, expression, localization, inferred interaction, genetic interactions, data mining, and neighborhood. Likewise, each node can be related to flags (events) related to the reported Hallmarks.

**Figure 3.**
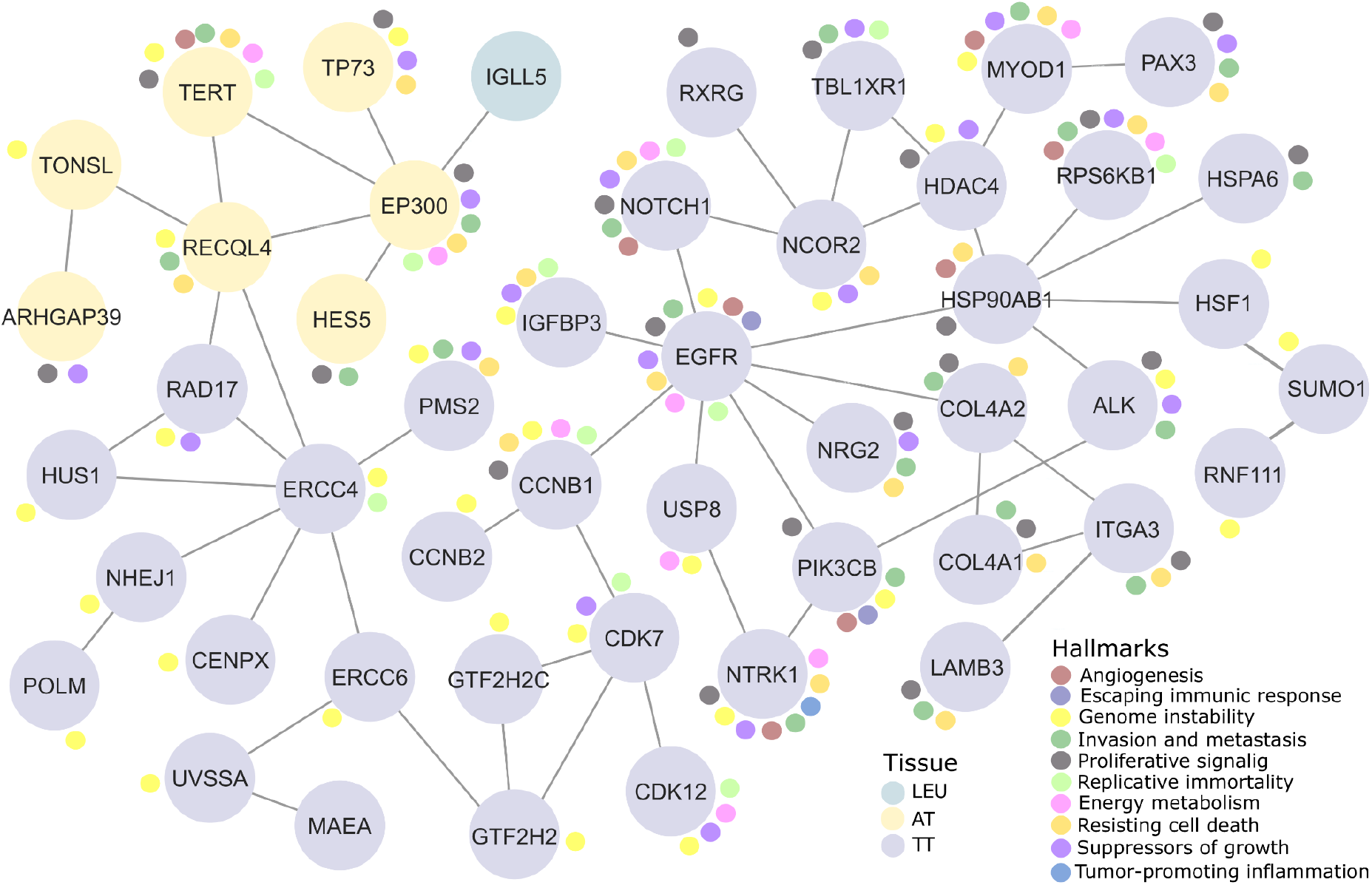
Relevant gene networks and the Hallmarks Cancer and metabolic pathways. Ten Hallmarks of Cancer are shown in the legend. The tumor tissues (TT) in purple, tumor adjacent tissues (AT) in yellow, and leukocytes (LEU) are indicated in blue light.

In this way, the network formed among LEU had one node with IGLL5. In the ADJ network, the genes with the highest number of nodes were EP300 and RECQL4. In the TUM network, there were 39 nodes; genes ERCC4, EGFR, and HSP90AB1 have high connections. EGFR, TERT, NTRK1, and RPS6KB1 have the most significant number of Hallmarks in the network (Table 6).

**Table 6.**
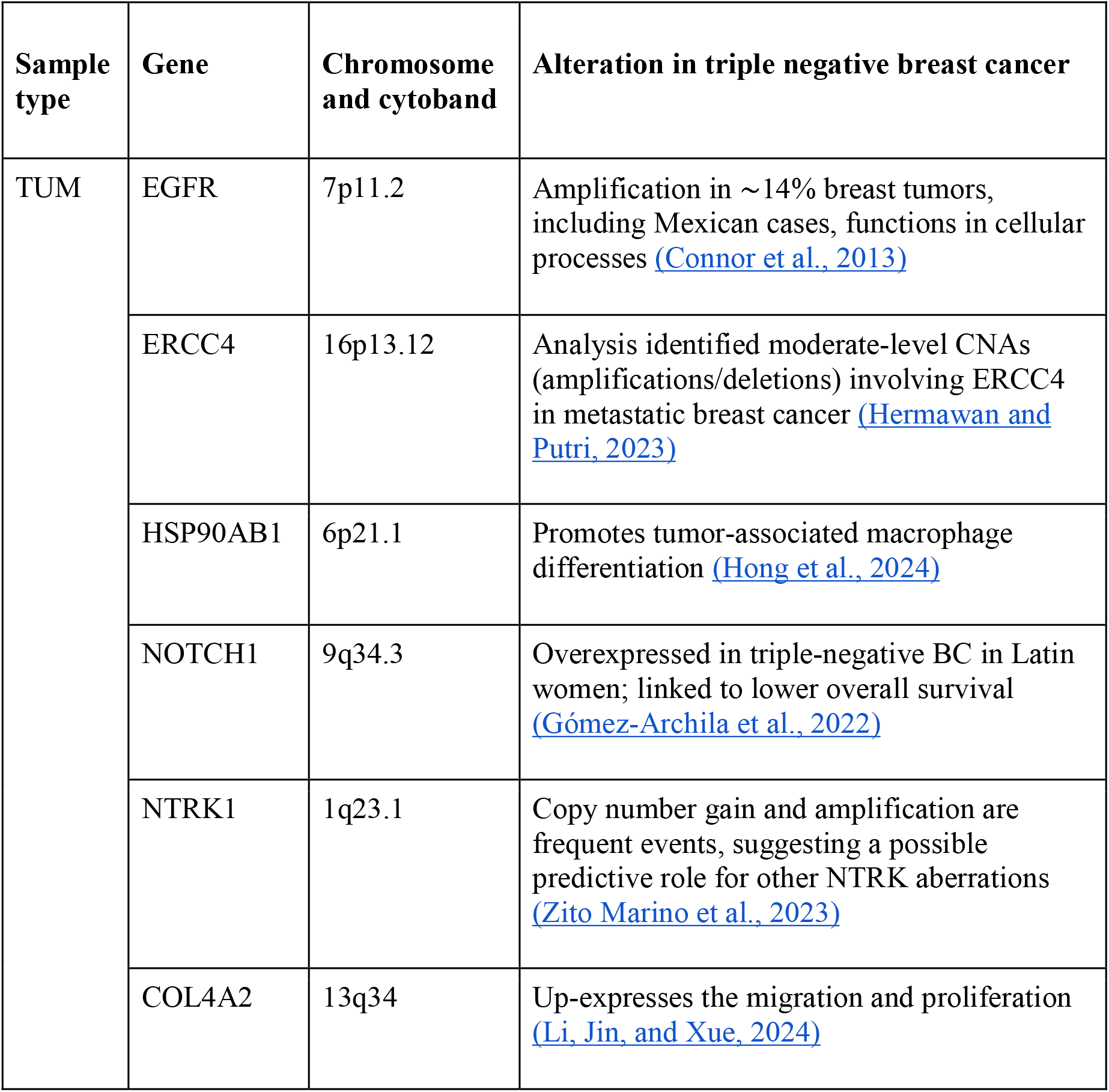
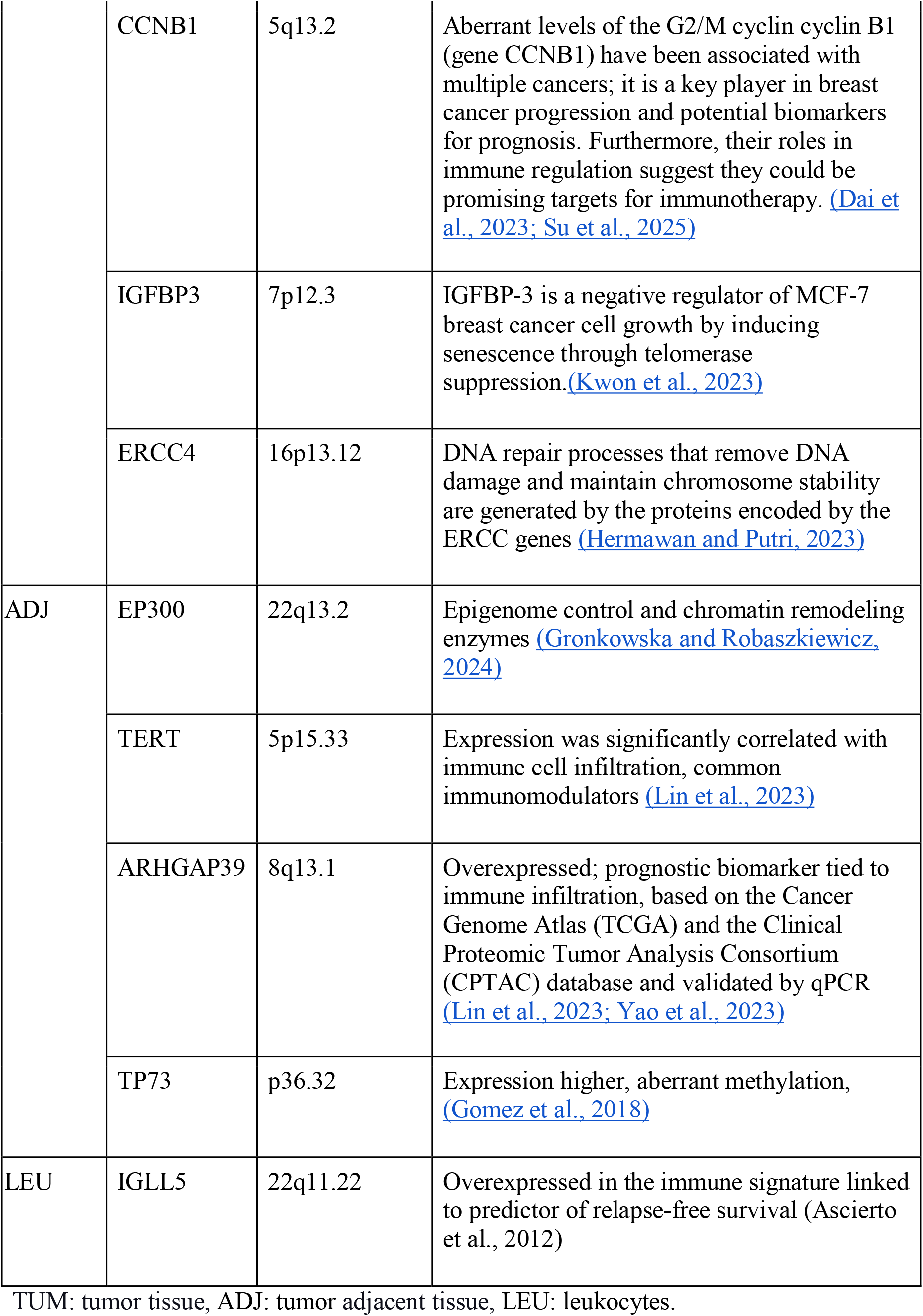
Top genes linked to the highest number of Hallmarks of Cancer and genes detected in the breast.

## Discussion

This study represents one of the first genome-wide high-density CNV analyses of TNBC in Mexican patients, including triads TUM, ADJ, and LEU. The results reveal a complex landscape of genomic alterations involving both shared and exclusive CNVs across these sample types. Our findings highlight the significance of CNVs not only in tumorigenesis but also in the tumor microenvironment and peripheral blood, suggesting a broader systemic genomic instability.

Chromosomal regions such as 1q23.3, 1q32.1, and 8q24.3 showed consistent gains across tumor samples. These loci harbor oncogenes like MCL1, BCL9, and MYC, which have been implicated in cancer progression, resistance to apoptosis, and metastasis (Curtis et al., 2012; Chial, 2008). In contrast, losses in 6q25.2—encompassing the ESR1 gene—are aligned with hormone-receptor negative phenotypes, reinforcing the clinical characteristics of TNBC. Furthermore, gains in 20q13.33 (ZNF217) and 8q24.3 were also detected in ADJ and LEU, supporting the notion that CNVs may originate early in tumorigenesis or reflect systemic alterations.

Hallmarks of Cancer enrichment analysis confirmed that tumor samples predominantly showed CNVs associated with processes such as proliferation, invasion, metastasis, and immune evasion, while ADJ samples were linked to growth suppression, and LEU samples to genomic instability. This stratification underlines the molecular heterogeneity of TNBC and offers a nuanced view of how genomic alterations influence tumor behavior and the microenvironment.

Functional pathway analysis further validated these findings. In TUM, pathways related to DNA repair (NER, TP53 signaling), extracellular matrix remodeling (collagen assembly), and cell cycle checkpoints (G2/M) were significantly enriched. These molecular disruptions are consistent with TNBC’s aggressive phenotype and its poor response to conventional therapies. Interestingly, ADJ showed enrichment in NOTCH and RAC3 signaling, suggesting that non-malignant cells within the tumor microenvironment may also undergo functional reprogramming, potentially contributing to tumor support or resistance mechanisms.

We also identified a subset of CNV-genes with high network centrality and multiple Hallmark associations—such as EGFR, ERCC4, and HSP90AB1—offering potential as markers or therapeutic targets. EGFR was highly connected within the network and amplified in several patients, consistent with its known role in TNBC and previous reports of overexpression in Latin American cohorts (Gómez-Archila et al., 2022; Connor et al., 2013).

An important aspect of our study is the inclusion of Mexican patients’ descent. Genomic studies often underrepresent Latin American populations, leading to limited generalizability of findings and complicating clinical interpretation (Fernández-López et al., 2019). Our results emphasize the importance of population-specific genomic studies to uncover potentially unique molecular signatures that may inform diagnosis, prognosis, or therapy in underrepresented groups.

Finally, while our results offer valuable insights, certain limitations must be acknowledged. The small sample size limits statistical power, and the lack of RNA expression data precludes functional validation of CNVs. Nonetheless, the integration of CNV profiles with Hallmark functions and interaction networks presents a robust framework for understanding TNBC heterogeneity and identifying candidate molecular targets.

We recognized the necessity of including populations from diverse ethnic backgrounds in genomic analyses. This inclusion is vital not only for increasing access to genetic testing but also for ensuring that the results of such tests are interpretable across different demographic groups. The current lack of representation poses a significant clinical challenge, known as a “double disparity,” which complicates the interpretation of genetic test results from individuals with non-European backgrounds (Fernández-Lopez et al., 2019).

Efforts were made to develop accurate and efficient methods for detecting genetic alterations, as CNVs can serve as crucial markers for tumors, with potential implications in tumor subtyping and predicting responses to therapies. However, reliable CNVs detection can be hindered by challenges such as normal cell contamination, tumor heterogeneity, and systematic errors resulting from structural variations. Therefore, it is recommended to identify these variations before conducting CNVs analysis on tumor samples.

## Conclusions

This study demonstrates the value of integrating high-density copy number variation (CNV) profiling with network-based analyses of tumor tissue, adjacent tissue, and blood samples. This approach offers a systemic view of genomic instability in triple-negative breast cancer (TNBC). Applying this approach to Mexican patients allowed us to identify recurrent alterations and central nodes in cancer-related pathways, while also contributing to reducing the underrepresentation of Latin American populations in genomic research. This methodological framework can be extended to other cohorts and cancer subtypes, facilitating the transition from descriptive variation maps to functionally relevant biomarkers. Notably, population-specific genomic references, like those presented here, lay the groundwork for the development of equitable precision oncology strategies and the fostering of collaborations that address global disparities in cancer research and care.

